# Dissecting the functional contributions of different neuronal subtypes in the dorsal striatum to perseverative behaviour in ephrin-A2/A5^-/-^ mice

**DOI:** 10.1101/2022.10.28.514217

**Authors:** Maitri Tomar, Jennifer Rodger, Jessica Moretti

## Abstract

Overreliance on habit is linked with disorders such as drug addiction and obsessive-compulsive disorder and there is increasing interest in the use of repetitive transcranial magnetic stimulation (rTMS) to alter neuronal activity in the relevant pathways and reduce relapse or accelerated shift towards habit formation. Here we studied the brains of ephrin-A2A5^-/-^ mice, which previously showed perseverative behaviour in progressive-ratio tasks, associated with low cellular activity in nucleus accumbens. We investigated if rTMS treatment had altered the hierarchical recruitment of brain regions from ventral to dorsal striatum associated with abnormal habit formation in these mice.

Brain sections of mice that underwent progressive-ratio tasks with and without low intensity rTMS (LI-rTMS) were taken from a previous study. We take advantage of the previous characterisation of perseverative behaviour to investigate the contribution of different neuronal subtypes and striatal regions. Striatal regions were stained for neuronal activation with c-Fos and for medium spiny neurons with DARPP32. Qualitative analysis was carried out for other neuronal subtypes in the striatum - GABAergic, parvalbumin-expressing and cholinergic interneurons.

Contrary to our hypothesis, we found neuronal activity in ephrin-A2A5^-/-^ mice still reflected goal-directed behaviour. However, we saw that the dorsolateral striatum contributed more to total striatal activity in untreated ephrin-A2/A5^-/-^ mice. This supported our hypothesis that ephrin-A2/A5^-/-^ mice have greater c-Fos activity in habit-associated striatal regions. LI-rTMS in ephrin-A2A5^-/-^ mice also appeared to delay the shift from goal-directed to habitual behaviour as suggested by increased activation in dorsomedial striatum and nucleus accumbens.

## 1 Introduction

Habits form over a period of time and require a shift from goal-directed behaviour to habitual behaviour. However, sometimes a faulty shift can contribute to disorders such as drug addiction (Everitt and Robbins, 2005) and obsessive-compulsive disorder (OCD) (Gillan et al., 2011). The formation of habit is generally defined as a shift from response-outcome associations, where actions are primarily mediated by the value associated with the outcome, to stimulus-response associations where, over a period of extended training and repetition, there is a pattern of outcome devaluation and the behaviour becomes more automatic, independent from the outcome of the action (Dickinson et al., 1995). This behavioural shift from goal-directed to habitual behaviour involves dopaminergic pathways and a hierarchical recruitment of brain regions from ventral to dorsal striatum (Segovia et al., 2012; Liu et al., 2013).

A potential tool to manipulate this hierarchical recruitment is repetitive transcranial magnetic stimulation (rTMS), a non-invasive brain stimulation technique that is able to modulate neuronal activity (Aydin-Abidin et al., 2008) and various neurotransmitters, including dopamine (Keck et al., 2002; Moretti et al., 2020). rTMS is approved for treatment of depression (O’Reardon et al., 2007) and OCD (Carmi et al., 2018). Understanding more about the mechanisms of rTMS, could be useful in refining rTMS use.

To explore the potential of rTMS to repair abnormal circuitry, a transgenic ephrin-A2/A5^-/-^ mouse strain has previously been used to measure structural and functional plasticity induced in abnormal neural pathways. Ephrins are membrane-bound ligands of Eph tyrosine kinase receptors and are important in cell migration and axon guidance during development (Wilkinson, 2001). As a result, ephrin-A2/A5^-/-^ mice, which lack the ephrin-A2 and ephrin-A5 ligands, have disorganized axonal projections in neuronal circuits throughout the brain due to disrupted Eph/ephrin signalling. These mice have abnormal visual topography and visuomotor behaviour which are both partially rescued by low-intensity rTMS (LI-rTMS) (Rodger et al., 2012; Makowiecki et al., 2014; Poh et al., 2018). In addition, the mice have reduced dopaminergic innervation of the striatum (Sieber et al., 2004; Cooper et al., 2009) associated with abnormal behavioural patterns in goal-directed behaviour (Poh et al., 2018) but the effects of LI-rTMS on this phenotype are less well understood.

A previous study (Moretti et al., 2021) explored whether the different goal-directed behaviour of ephrin-A2/A5^-/-^ mice compared to wild type was due to abnormal motivation by comparing performance of both strains in a progressive ratio (PR) tasks. PR tasks are a good measure of motivation as they involve an increasing instrumental response requirement for a reward. Highly motivated animals will continue to respond consistently, while animals with low motivation stop or slow their responding (Hodos, 1961; Aberman et al., 1998). The potential ameliorating effect of chronic excitatory LI-rTMS delivered during the PR task was also examined. Unexpectedly, the results did not support a motivation phenotype: ephrin A2/A5^-/-^ mice showed perseverative behaviour in the PR task, with a greater number of responses compared to wildtype mice despite increasing task difficulty (Moretti et al., 2021). Compared to wildtype mice, ephrin-A2A5^-/-^ mice also showed reduced c-Fos expression in the nucleus accumbens (NAc) after two weeks of the PR task (Moretti et al., 2021), which may reflect an accelerated dorsal shift in neuronal activity characteristic of habitual behaviour (Segovia et al., 2012). Interestingly, chronic excitatory LI-rTMS delivered to ephrin-A2/A5^-/-^ mice during the PR task did not significantly alter behaviour, but did ameliorate the reduced c-Fos expression in the ventral striatum of ephrin-A2/A5^-/-^ mice, resulting in expression that was similar to wildtype levels (Moretti et al., 2021). Therefore, in ephrin-A2/A5^-/-^ mice, LI-rTMS may mitigate abnormal activity along the mesolimbic pathway associated with behavioural inflexibility, by delaying an accelerated shift in neuronal activity toward the dorsal striatum.

To understand if altered dorsal striatum activity in ephrin-A2/A5^-/-^ mice was responsible for perseverative behaviour in these mice and if rTMS could delay an accelerated shift from goal-directed behaviour to habit, here we used tissue from the previous study (Moretti et al., 2021) to identify and compare activation of neuronal populations across the entire striatum, investigating the dorsomedial striatum (DMS), dorsolateral striatum (DLS) and NAc (core and shell) using c-Fos immunohistochemistry. We compared c-Fos expression in different regions of the striatum in wildtype and ephrin-A2A5^-/-^ mice that either received active LI-rTMS or sham stimulation. In addition, since a low percentage of c-Fos expression colocalised with medium spiny neurons (MSNs) was found in the NAc (Moretti et al., 2021), we also identified the other neuronal subtypes in the striatum that expressed c-Fos including GABAergic and cholinergic interneurons. Our in depth characterisation of cell activation following LI-rTMS in the striatum in behaviourally characterised animals may provide insights into the use of TMS for disorders such as addiction and OCD where the shift of habit formation is faulty.

## 2 Materials and methods

### 2.1 Animal tissue

All experiments were approved by The University of Western Australia Animal Ethics Committee (AEC 100/1639). Sagittal mouse brain sections (40 µm) from 11 adult (8-24-week-old) wildtype C57BL6/J mice (sham: 6; rTMS: 5) and 10 ephrin-A2/A5^-/-^ mice (sham: 6; rTMS: 4) (Moretti et al., 2021) were used. The ephrin-A2/A5^-/-^ mice was backcrossed onto C57BL6/J mice for >20 generations, bread from heterozygous ephrin-A2^-/-^A5^+/-^ parents and genotyped at weaning (Feldheim et al., 2000). In the previous study, following habituation to the LI-rTMS coil and training in the operant box, mice performed one week of an exponential PR task (required responses increase under an exponential equation) followed by one week of PR7 (requirement started at 7 responses and increased by 7 with each reinforcement) (Moretti et al., 2021). Mice received 14 days of biomimetic high frequency stimulation (BHFS) LI-rTMS or sham during the first 10 minutes of the daily PR task and were euthanized 90 minutes after the beginning of the final PR task to correspond with the peak of c-Fos expression (Moretti et al., 2021). A summary of the experimental design from Moretti et al. (2021) is reproduced in Figure 1. The previous study showed perseverative behaviour in untreated ephrin-A2/A5^-/-^ mice and low c-Fos expression in the NAc which improved following LI-rTMS (Moretti et al., 2021). This suggested an accelerated shift from goal-directed to habit in untreated ephrin-A2/A5^-/-^ mice which could have been delayed by LI-rTMS (Moretti et al., 2021). However, in the previous study neuronal activation in dorsal striatum was not investigated. In the current study we use additional tissue from the same animals as Moretti et al. (2021) to confirm the shift in neuronal activity from regions involved in goal-directed behaviour to habit and if LI-rTMS delayed it on a cellular level.

**Figure 1.**
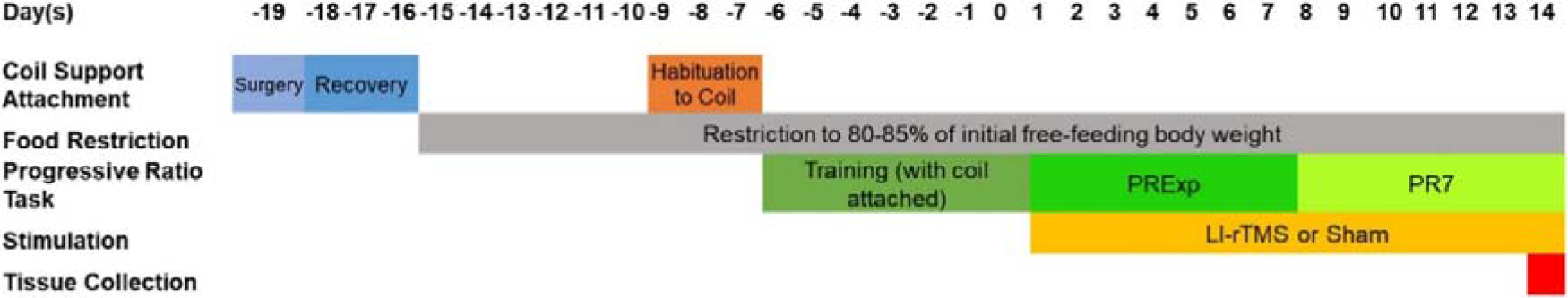
Outline of the experimental paradigm performed by Moretti et al. (2021). Reproduced from (Moretti et al., 2021) with permission from Behavioural Brain Research.

### 2.2 Immunofluorescence

Every fifth sagittal brain section which contained dorsal striatum was identified using the mouse brain atlas (Paxinos and Franklin, 2012). These sections were stained for three different antibodies: c-Fos (rabbit polyclonal c-Fos antibody, 1:5000, Abcam, ab190289), a marker for neuronal activity (Bullitt, 1990), cAMP-regulated phosphoprotein-32 kD (DARPP32) (purified mouse anti-DARPP32, 1:2000, BD Transduction Laboratories, 611520), a marker for MSNs (Anderson and Reiner, 1991), and glutamate decarboxylase 67 (GAD67) (anti-Goat, 1:750, R&D systems, AF2086-SP), a marker for GABAergic neurons (Lazarus et al., 2015). For confirmatory analysis of active parvalbumin (PV)-expressing GABAergic interneurons and cholinergic interneurons, two sections from one animal belonging to each experimental group were stained for c-Fos, PV (mouse monoclonal anti-PV antibody, 1:5000, Millipore Sigma, p3088) and ChAT (anti-choline acetyltransferase antibody, 1:200, Millipore Sigma, AB144P). Hoescht (1:2000, Invitrogen) was used to stain all nuclei.

Free-floating sections were washed 3x in PBS (5 min each), permeabilised by washing in 0.1% Triton-X in PBS (PBS-T) for 15 min and incubated for 2 h in a blocking buffer of 2% bovine serum albumin (Sigma) and 3% normal donkey serum (Sigma) diluted in 0.1% PBS-T. Then, sections were incubated with primary antibodies in blocking buffer overnight at 4°C. Sections were washed in PBS-T 3x (10 min each) and incubated for 2 h with secondary antibodies diluted in blocking buffer to 1:600 (donkey anti-rabbit IgG Alexa Fluor 488, Invitrogen, Thermo Fisher, A21206; donkey anti-mouse IgG Alexa Fluor 555, Invitrogen, Thermo Fisher, A21202; donkey anti-goat IgG H&L Alexa Fluor 647, ab150131). Sections were then washed in PBS-T 3x (10 min each) and incubated for 10 min in Hoescht in PBS (1:1000, Invitrogen). Lastly, the sections were washed in PBS 3x (10 min each) and mounted on gelatin-subbed slides, coverslipped with mounting medium (Dako, Glostrup, Denmark) and sealed with nail polish. Slides were stored at 4°C in a light-controlled environment until imaging.

### 2.3 Imaging and quantification

Commonly used mouse brain atlases (Lein et al., 2007; Paxinos and Franklin, 2012) do not show detailed segmentation of the dorsal striatum (Chon et al., 2019). Therefore, in the present study the two regions of the dorsal striatum (DMS and DLS) were identified by creating a custom sagittal atlas that combined the location information of dorsal striatum sections from a mouse brain atlas (Chon et al., 2019) annotated onto a 3D magnetic resonance imaging (MRI) data of P56 mouse brain atlas using ITK-SNAP (Yushkevich et al., 2006) software (Fig 2).

**Fig 2.**
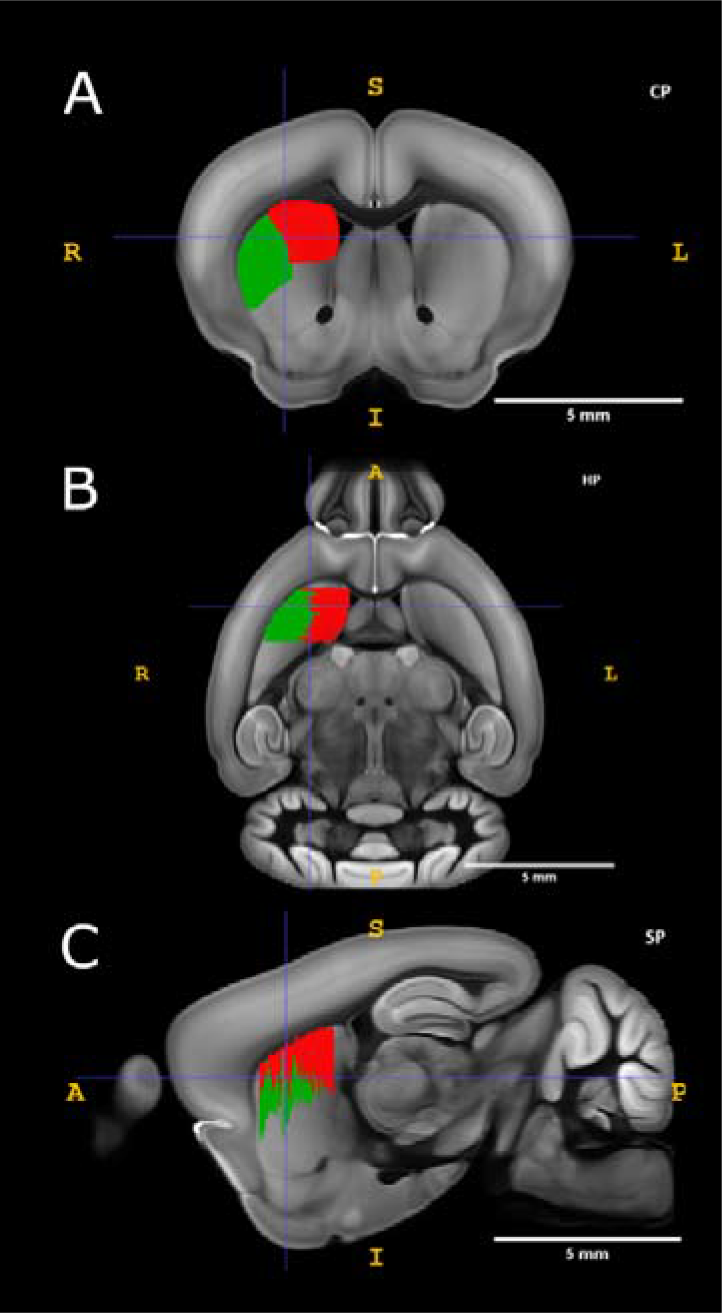
Identification of regions of interest on a mouse brain atlas. (A) Location of DMS and DLS was marked in red and green respectively on a coronal plane in the software, ITK-SNAP which produced corresponding images in the horizontal (B) and sagittal planes (C). Scale bar = 5 mm. CP, coronal plane; SP, sagittal plane; HP, horizontal plane.

Sections were imaged through z-stacks (2 µm apart) on a Nikon C2 confocal microscope (Nikon, Tokyo, Japan). 10x magnification was used to identify the regions of interest. 8-11 fields of view (FOV) for DLS and 6-10 FOV for DMS across 5-7 sections each animal were used to image at 40x magnification to identify c-Fos, DARPP32 and GAD67 markers. For the confirmatory analysis, 2 FOV for DLS and DMS and 1 FOV for NAc across 2 sections from each experimental group was imaged at 40x magnification.

Cell counts were performed using stereological principles using ImageJ software and were carried out blinded to experimental group after all images were captured. The total number of c-Fos single labelled cells, c-Fos^+^/DARPP32^+^ double labelled cells, c-Fos^+^/GAD67^+^/DARPP32^-^ double labelled cells, c-Fos^+^/PV^+^ double labelled cells and c-Fos^+^/ChAT^+^ double labelled cells were identified and counted for each image (Fig 3). The combination of DARPP32^+^ and GAD67^+^ double labelling was not assessed since DARPP32^+^ cells are also GAD67^+^ because they are GABAergic in nature. c-Fos^+^ cells not colocalized with DARPP32 were classified as unspecified c-Fos.

**Fig 3.**
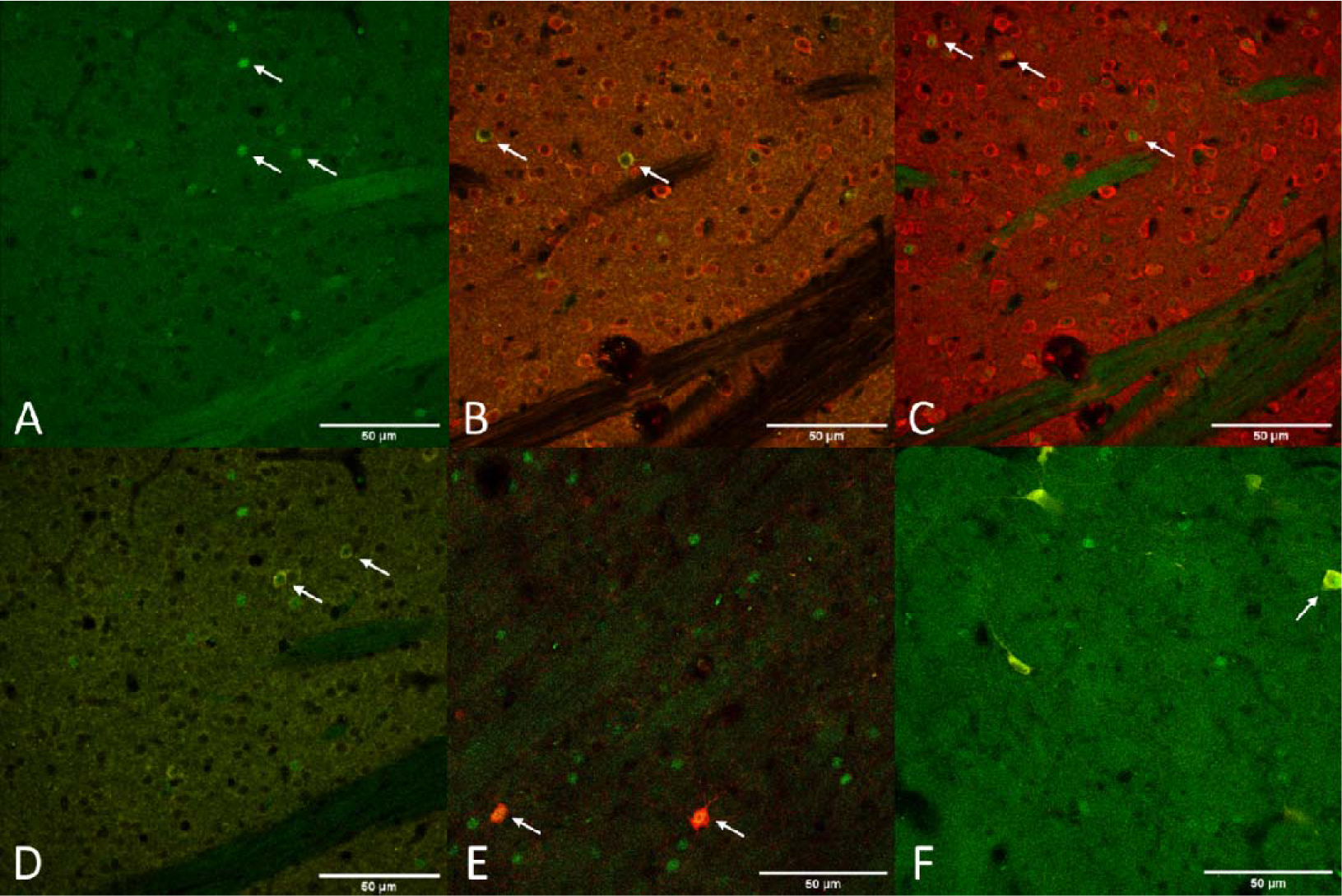
Representative images taken at 40x magnification showing different marker expressions. (A) expression of c-Fos^+^ cells (green) (B) expression of GAD67^+^ cells (yellow) and DARPP32^+^ cells (red) (C) expression of c-Fos^+^ (green) /DARPP32^+^ (red) colocalised cells (D) expression of c-Fos^+^ (green) /GAD67^+^ (yellow) colocalised cells (E) expression of c-Fos^+^ (green) /ChAT^+^ (yellow) colocalised cells (F) expression of c-Fos^+^ (green) /PV^+^ (red) colocalised cells. Scale bar = 50 µm. GAD67, Glutamate decarboxylase 67; cAMP-regulated phosphoprotein-32 kD, DARPP32.

For confirmatory analysis, c-Fos-labelled cells colocalised with GAD67, PV and ChAT were counted and analysed to explain unspecified c-Fos counts. During imaging, sections from one ephrin-A2/A5^-/-^-rTMS animal was excluded due to poor staining (ephrin-A2/A5^-/-^-rTMS *n = 3*).

### 2.4 Statistical analysis

For analysis, the total c-Fos cells/mm^3^ and percentage of c-Fos cells colocalised with DARPP32, GAD67, PV and ChAT were calculated for each animal. Statistical analysis was not done on counts showing c-Fos colocalization with PV and ChAT due to low sample size but assessed qualitatively.

Assumptions of normality and homogeneity were checked. For DLS counts, the assumptions of normality were not met for percentage c-Fos-GAD67 and total c-Fos cells/mm^3^ counts (p<0.05). Homogeneity of variance was checked, and it was met for all data sets (Levene’s test, p>0.05). For DMS counts, the assumptions of normality were not met for percentage c-Fos-DARPP32, percentage c-Fos-GAD67 and total c-Fos cells/mm^3^ (p<0.05). The assumption of homogeneity of variance was not met for percentage c-Fos-DARPP32 data (Levene’s test, p<0.05) but corrected with logarithmic transformation.

To compare whether cell counts changed across groups, multiple two-way analysis of variance (ANOVAs) were used with the independent variables of strain and treatment. Each ANOVA had the dependent variable of either total c-Fos cells/mm^3^, percentage c-Fos-DARPP32 or percentage unspecified c-Fos cell count data and were run separately for both DLS and DMS. ANOVA was also performed for c-Fos^+^/GAD67^+^/DARPP32^-^ cell counts for the DLS region. For the DMS regions, due to small sample size, Ephrin-rTMS group was excluded from statistical analysis for c-Fos^+^/GAD67^+^/DARPP32^-^ data set and student t-tests were performed to compare remaining experimental groups.

Additional data for percentage of c-Fos^+^/GAD67^+^/DARPP32^-^ cell counts were also obtained for NAc core and shell if the regions were present in the stained tissue (NAc core, Group: wildtype-sham *n=5,* wildtype-rTMS *n=4,* ephrin-A2/A5^-/-^-sham *n=4,* ephrin-A2/A5^-/-^-rTMS *n=*2; NAc shell, Group: wildtype-sham *n=6,* wildtype-rTMS *n=5,* ephrin-A2/A5^-/-^-sham *n=4,* ephrin-A2/A5^-/-^-rTMS *n=*3). This was combined with total c-Fos cells/mm^3^ and percentage c-Fos-DARPP32 cell counts for NAc core and shell from the previous study (Moretti et al., 2021). The percentage of c-Fos-GAD67 data set in NAc core and shell assumptions of normality and heterogeneity. Therefore, non-parametric Mann-Whitney U tests were performed to compare the percentage of c-Fos^+^/GAD67^+^/DARPP32^-^ cell counts between sham and treatment for both wildtype and ephrin-A2A5^-/-^ groups.

## 3 Results

### 3.1 Total c-Fos cells/mm^3^

In DLS, there was no overall effect of strain (F(1,16)=1.040, p=0.323, η^2^=0.058) or treatment (F(1,16)=0.973, p=0.339, η^2^=0.054) on c-Fos cells/mm^3^. The strain*treatment interaction was also not significant (F(1,16)=0.0007, p=0.979, η^2^=0.00) (Fig 4). Similarly, in DMS there was no significant difference in cell counts between strain (F(1,16)=0.902, p=0.356, η^2^=0.046) and no overall effect of treatment was observed (F(1,16)=1.342, p=0.264, η^2^=0.068). No significant strain*treatment interaction was observed (F(1,16)=1.378, p=0.258, η^2^=0.070) (Fig 4).

**Fig 4.**
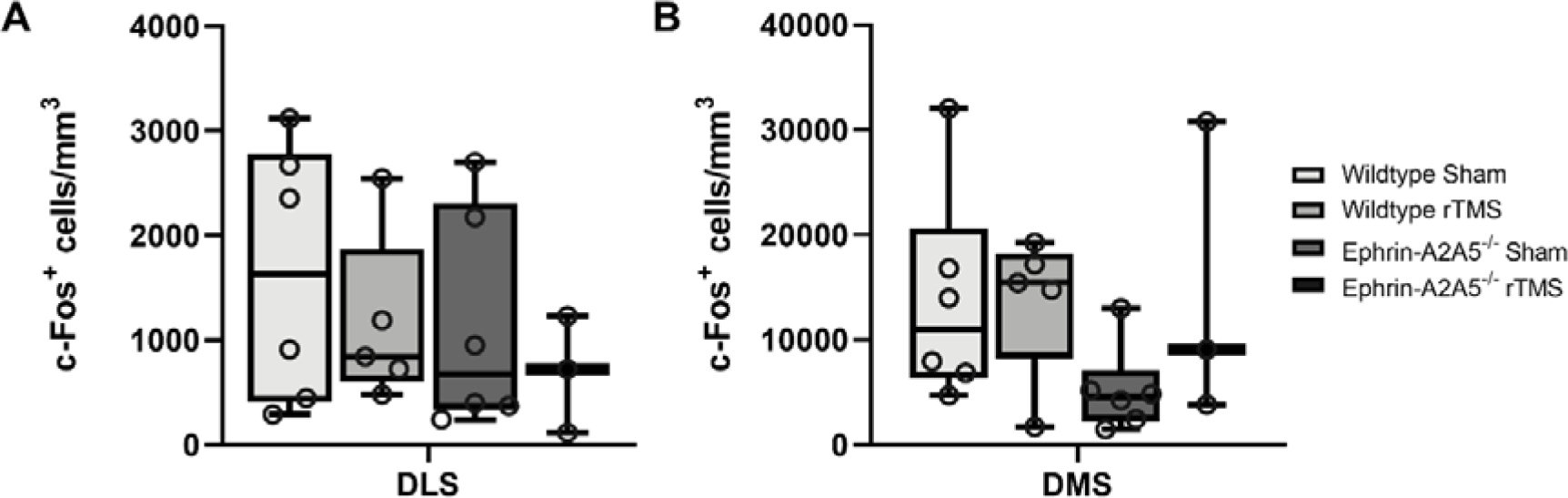
Graphs showing total c-Fos^+^ cells/mm^3^ counts for each group in (A) DLS and (B) DMS. No significant difference in total c-Fos**^+^** cells/mm^3^ cells was found between groups in both DLS and DMS (p>0.05). Points represent individual animal counts. Error bars represent ± SEM.

### 3.2 c-Fos colocalization in MSNs

In DLS, there was no overall effect of strain (F(1,16)=0.100, p=0.756, η^2^=0.006) or treatment (F(1,16)=0.001, p=0.976, η^2^=0) on percentage of c-Fos-DARPP32 colocalization. The strain*treatment interaction was also not significant (F(1,16)=0.376, p=0.548, η^2^=0.023) (Fig 5). Similarly, in DMS there was no significant difference in percentage of colocalization between strains (F(1,16)=0.0461, p=0.833, η^2^=0.002) and no overall effect of treatment was observed (F(1,16)=1.2556, p=0.279, ^2^=0.056). However, there was a significant strain*treatment interaction (F(1,16)=5.1062, p=0.038, η^2^=0.228). To understand the interaction, we followed up with pairwise post-hoc comparisons using Tukey’s test. There was an increase in percentage of c-Fos-DARPP32 colocalization after treatment in ephrin-A2/A5^-/-^ mice compared to ephrin-sham group (Fig 5), however the effect did not survive multiple comparison Tukey correction (uncorrected p=0.041, t(16)=-2.225, Tukey corrected p=0.159) despite the significant omnibus interaction. All other groups were non-significantly different from each other.

**Fig 5.**
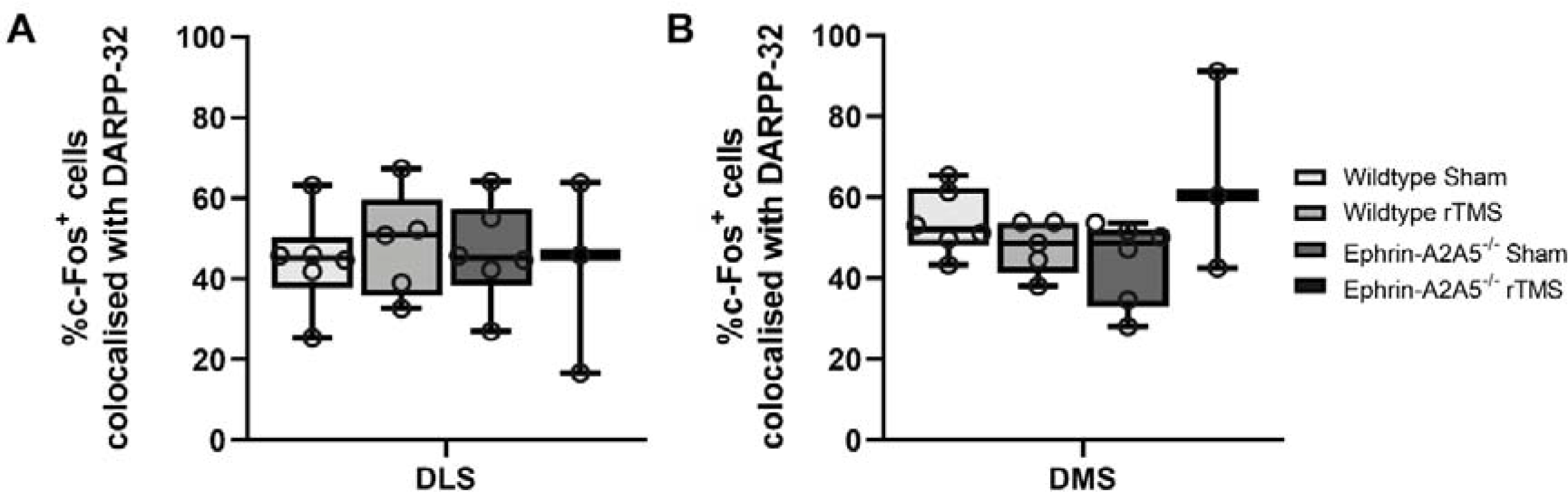
Graphs showing percentage of c-Fos^+^ cells colocalized with DARPP32^+^ cells for each group in (A) DLS and (B) DMS. Following stimulation, a significant increase was only seen in c-Fos-DARPP32 colocalization in DMS in ephrin-A2/A5^-/-^ mice in the omnibus test but not in pairwise post-hoc comparisons. Points represent individual animal counts.Error bars represent ± SEM.

### 3.3 Percentage of unspecified c-Fos cells

In DLS, there was no overall effect of strain (F(1,16)=1.9052, p=0.186, η^2^=0.105) or treatment (F(1,16)=0.0309, p=0.863, η^2^=0.002) on the percentage of unspecified c-Fos cells. The strain*treatment interaction was also not significant (F(1,16)=0.2026, p=0.659, η^2^=0.011) (Fig 6). Similarly in DMS, there was no overall effect of strain (F(1,16)=0.482, p=0.497, η^2^=0.026) or treatment (F(1,16)=0.270, p=0.611, ^2^=0.015) on the percentage of unspecified c-Fos cells. The strain*treatment interaction was also not significant (F(1,16)=1.738, p=0.206, η^2^=0.094) (Fig 6).

**Fig 6.**
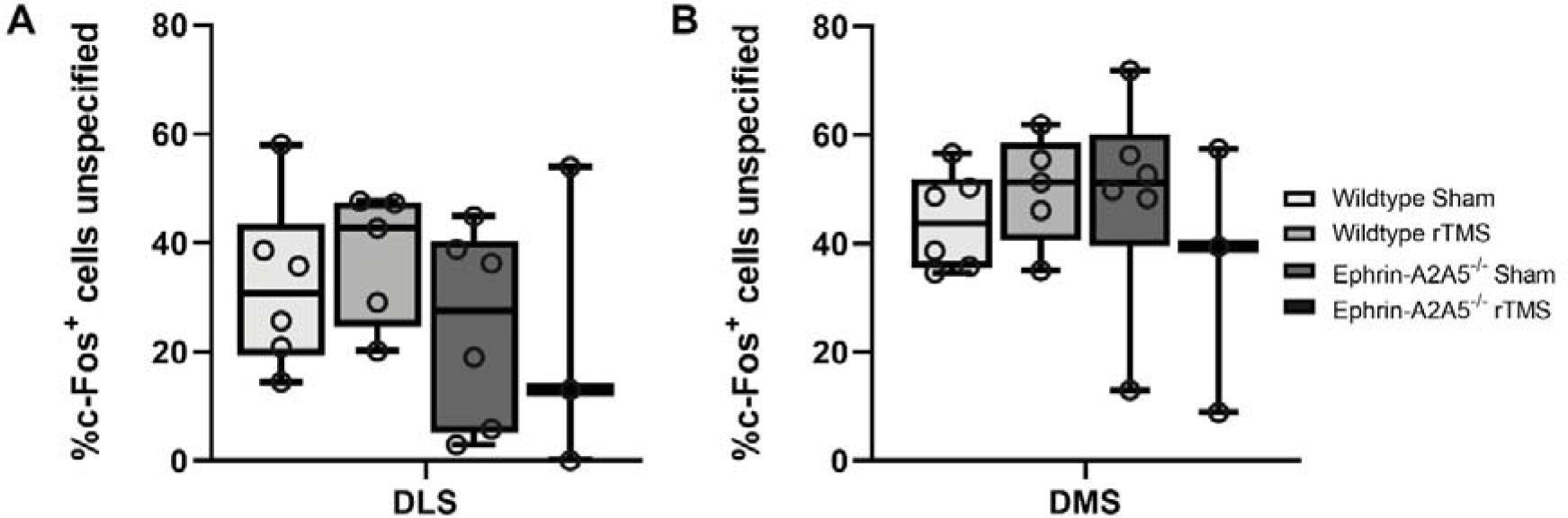
Graphs showing percentage of unspecified c-Fos^+^ cells for each group in (A) DLS and (B) DMS. No significant difference in percentage of c-Fos**^+^** cells that are single-labelled (c-Fos^+^/GAD67^-^/DARPP32^-^) was found between groups in both DLS and DMS (p>0.05). Points represent individual animal counts. Error bars represent ± SEM.

#### 3.3.1 Confirmatory analysis for unspecified c-Fos-labelled cells

Across all experimental groups, approximately 60% and 40% of c-Fos labelled cells in the ventral and dorsal striatum, respectively were unidentified (i.e., not MSNs). Therefore, the sections were first stained for GABAergic interneurons to identify unspecified c-Fos labelled cells. Qualitative observation suggests that in DMS there were more active GABAergic interneurons as compared to DLS (Fig 7) and NAc where the percentage of colocalization across all groups were less than 1% and 2%, respectively.

**Fig 7.**
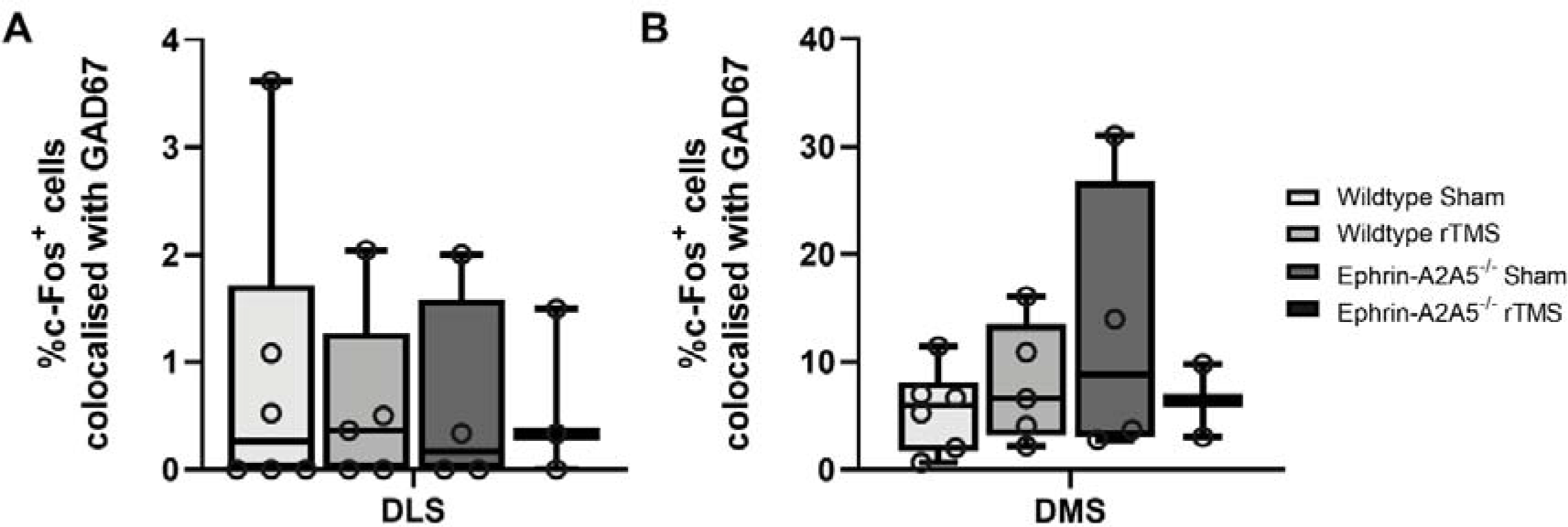
Graphs showing percentage of c-Fos^+^ cells colocalized with GAD67^+^-DARPP32^-^ cells for each group in (A) DLS and (B) DMS. No significant difference in percentage of c-Fos^+^ cells colocalised with GAD67^+^-DARPP32^-^ cells were found between groups in both DLS and DMS (p>0.05). Points represent individual animal counts. Error bars represent ± SEM.

For percentage of c-Fos^+^/GAD67^+^/DARPP32^-^ colocalization in DLS, no overall effect of strain (F(1,14)=0.058, p=0.814, η^2^=0.004) or treatment (F(1,14)=0.061, p=0.809, η^2^=0.004) was observed. The strain*treatment interaction was also not significant (F(1,14)=0.088, p=0.771, η^2^=0.006) (Fig 7). In the DMS, there was no significant change in c-Fos^+^/GAD67^+^/DARPP32^-^ colocalization between Wildtype-Sham and Wildtype-rTMS mice (t(9)=-0.864, p=0.410), and there was no significant difference between strains, Wildtype-Sham and Ephrin-Sham (t(8)=-1.32, p=0.222). Qualitatively, c-Fos^+^ that is colocalised with GAD67^+^decreased following LI-rTMS in ephrin-A2A5^-/-^ mice as seen in Fig 7.

In NAc core, there was no significant difference in the percentage of c-Fos^+^/GAD67^+^/DARPP32^-^ colocalization following treatment in both wildtype (U=8.5, p=0.771) and ephrin-A2A5^-/-^ mice (U=2, p=0.289). For the same data set, there was no significant difference between strains, Wildtype-Sham and Ephrin-Sham (U=6, P=0.240). Similarly, in NAc shell there was no significant difference in the percentage of c-Fos^+^/GAD67^+^/DARPP32^-^ colocalization following treatment in both wildtype (U=12, p=0.597) and ephrin-A2A5^-/-^ mice (U=5, p=0.825). There was also no significant strain difference, Wildtype-Sham and Ephrin-Sham (U=11, p=0.896).

As the percentage of c-Fos colocalised with GAD67-labelled cells were also low in both dorsal and ventral striatum, additional confirmatory tests were done for PV-expressing GABAergic interneurons and cholinergic interneurons. Qualitative analysis suggests there were more active PV-expressing GABAergic interneurons in DMS as compared to DLS and NAc. In DMS, a slight increase in active PV-expressing GABAergic interneurons was observed in ephrin-A2A5^-/-^ mice following treatment, whereas a slight decrease was seen in wildtype mice following treatment. No active cholinergic interneurons were observed in both DMS and NAc across all groups. In DLS, only about 2% c-Fos^+^ cells were colocalised with cholinergic interneurons in the wildtype-sham group, in all other groups no active cholinergic interneurons were observed.

### 3.4 Comparison of c-Fos activity across regions and groups

To further explore a potential change in activity distribution across regions in the overall striatum, we examined whether the proportion of c-Fos activity per regions differed between groups. We ran a 4*4 contingency χ^2^ test with c-Fos counts per region (NAc shell, NAc core, DMS and DLS) compared across experimental groups. The initial 4*4 contingency χ^2^ test was significant (χ^2^ (9, N=666524) =8030, p<0.001, V=0.0634). We then ran follow up 2*4 contingency χ^2^ test to understand where the difference was, comparing the proportion of striatal c-Fos activity per region for each group with the Wildtype-Sham group. Proportions of striatal c-Fos activity per region differed in comparison with Wildtype-Sham for all groups, although the test may be overly sensitive due to the large sample size (Wildtype-rTMS: χ^2^(3, N=436329) =642, p<0.001, V = .0384); Ephrin-Sham: χ^2^(3, N=335837) = 2156, p<0.001, V = .0801; Ephrin-rTMS: χ^2^ (3, N=384288) =2446, p<0.001, V = .0798). Although there are significant differences, the effect sizes for each comparison are small.

To further understand where the difference in striatal activity per region occurred, we performed region-by-region comparison for the percentage total striatal c-Fos activity present for each region across groups (Fig 8). Comparison between each experimental group and Wildtype-Sham used multiple Mann-Whitney U tests. Statistics are presented in Table 1. Despite significant comparisons on the group level, no regions appeared to be statistically different, however some effect sizes when comparing ephrin-A2A5^-/-^ groups with Wildtype-Sham were moderate (0.3-0.5) suggesting that the proportion of c-Fos in several regions for ephrin-A2/A5^-/-^ mice tended to be larger than Wildtype-Sham.

**Fig 8.**
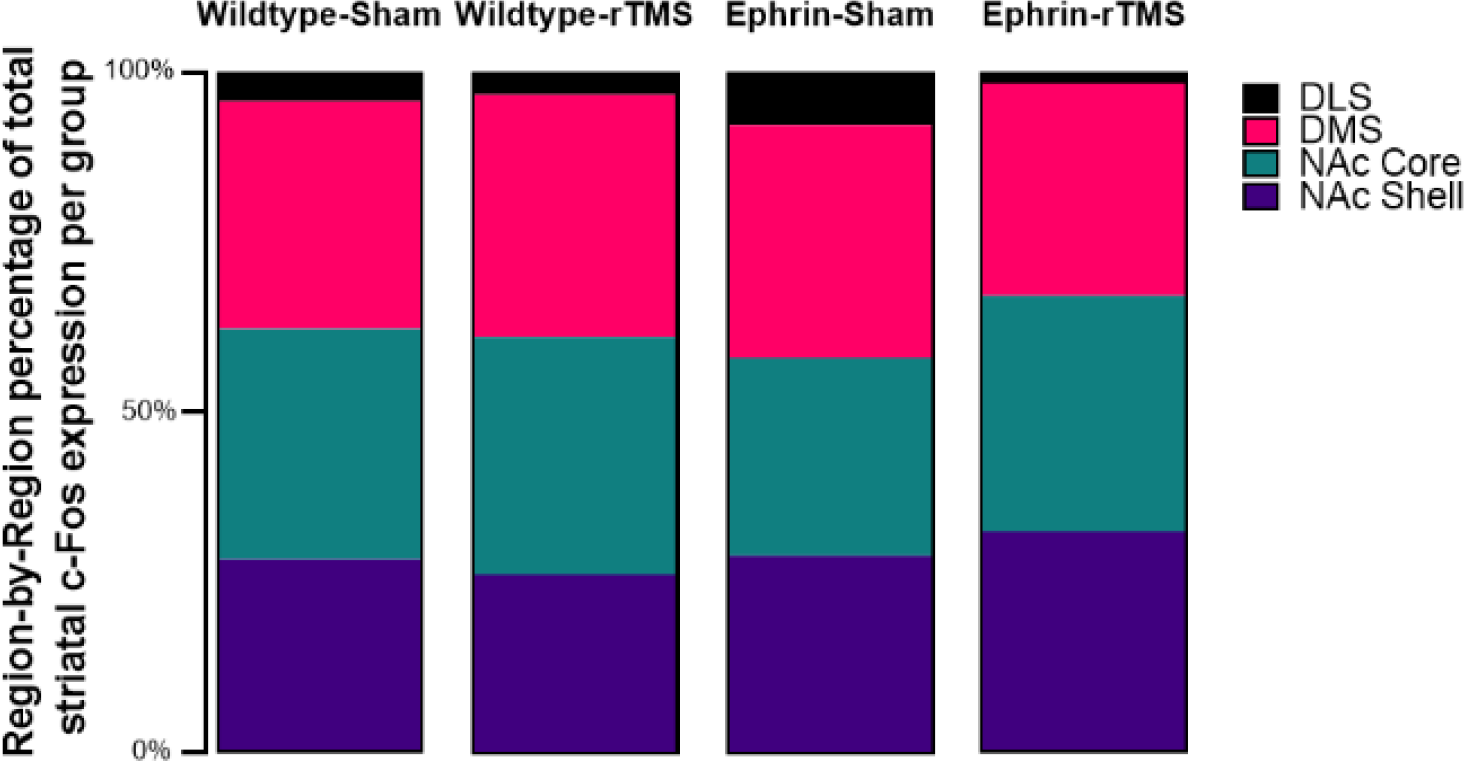
Average region-by-region proportion of total striatal activity in the experimental groups.

**Table 1.** Region by region Mann-Whitney U comparisons for proportion of striatal c-Fos.

| Group comparison<br>(Wildtype-Sham vs.) | Mann-Whitney<br>U | p | Effect Size<br>(Rank<br>biserial<br>correlation) | Mean<br>Difference | 95% Confidence<br>Interval |  |
| --- | --- | --- | --- | --- | --- | --- |
|  |  |  |  |  | Lower | Upper |
| **Wildtype-rTMS** |  |  |  |  |  |  |
| ***Region*** |  |  |  |  |  |  |
| DLS | 14 | 0.926 | 0.0667 | 0.271 | -2 | 4 |
| DMS | 11 | 0.537 | 0.267 | -5.5 | -22 | 21 |
| NAc Core | 14.5 | 1 | 0.0333 | 0.00368 | -18 | 14 |
| NAc Shell | 12 | 0.662 | 0.200 | 5.5 | -11 | 17 |
| **Ephrin-Sham** |  |  |  |  |  |  |
| ***Region*** |  |  |  |  |  |  |
| DLS | 11.5 | 0.333 | 0.361 | -3.88 | -21 | 3 |
| DMS | 15 | 0.699 | 0.167 | -5 | -24 | 20 |
| NAc Core | 11 | 0.31 | 0.389 | 9.5 | -9 | 27 |
| NAc Shell | 17 | 0.936 | 0.056 | 3 | -13 | 13 |
| **Ephrin-rTMS** |  |  |  |  |  |  |
| ***Region*** |  |  |  |  |  |  |
| DLS | 4.5 | 0.276 | 0.500 | 2 | -2 | 6 |
| DMS | 7 | 0.714 | 0.222 | 4.5 | -20 | 42 |
| NAc Core | 9 | 1 | 0 | -0.0727 | -23 | 17 |
| NAc Shell | 6 | 0.548 | 0.333 | -4.5 | -21 | 8 |

## 4. Discussion

Evidence of a habitual behavioural responding pattern in ephrin-A2/A5^-/-^ mice led to the hypothesis that these mice would display a shift in neuronal activity towards striatal regions involved in habit formation (Moretti et al., 2021). However, contrary to the hypothesis, neuronal activation was mostly seen in the regions of the striatum that are involved in goal-directed behaviour. We also hypothesized that the LI-rTMS change in c-Fos densities seen previously in ephrin-A2/A5^-/-^ mice NAc would extend to the dorsal striatum. In partial support of the hypothesis, we did not observe a change in total c-Fos expression after LI-rTMS in ephrin-A2/A5^-/-^ mice, but in the DMS there was an increase in MSN activation following stimulation. Although the proportion of c-Fos labelling across the striatum was significantly different in all experimental groups compared to control animals, there was not one specific region driving this difference.

### 4.1 Neuronal activation assessed by c-Fos expression was highest in regions involved in goal-directed behaviour

The overall proportion of c-Fos + cells was significantly different between Wildtype-Sham group and all other groups. However, comparing the difference in proportions across regions did not identify specific regions that were significantly different to the Wildtype-Sham group. Nonetheless, the biggest effect sizes were associated with the difference between Wildtype-Sham and the Ephrin groups, suggesting less activity in the DLS of Ephrin-rTMS while the comparisons with Ephrin-Sham animals suggest less activity in the NAc and greater activity in the DLS compared to Wildtype-Sham animals. Therefore, it appears that there is an increased proportion of dorsal striatum activity in Ephrin-Sham animals compared to Wildtype animals, and this difference is reduced following LI-rTMS. This is in line with our hypothesis that LI-rTMS may delay an accelerated shift from goal-directed behaviour to habit seen in ephrin-A2/A5^-/-^ mice.

While looking at the change in c-Fos activity in specific striatal regions, it was found that in both sham and treatment groups of ephrin-A2/A5^-/-^ mice, DMS had more c-Fos expression than DLS. Although DMS is a part of the nigrostriatal pathway, it is a transition zone and is versatile in its function as it is involved in both stimulus-response and action-outcome learning (Corbit et al., 2001) Therefore, DMS shows more activity when the behaviour is goal-directed, and this activity slowly decreases with over-training and shifts to DLS for formation of habits (Lipton et al., 2019). Similarly, there is a progression of c-Fos expression during training in a fixed-ratio (FR) task, from higher c-Fos activity in NAc shell on the first day of the task, towards the NAc core, DMS and finally to DLS as training progressed (Segovia et al., 2012). In our study, the major proportion ofc-Fos+ cells remained in the NAc core and DMS, suggesting that behaviour in all mice remained goal-directed (Fig 8). However, the Ephrin-Sham group showed signs of higher relative DLS activity, and less NAc core activity compared to Wildtype-Sham group which supports our hypothesis that behaviour may have shifted towards habit formation in ephrin-A2/A5^-/-^ mice. A caveat is that, unlike previous studies (Segovia et al., 2012), using c-Fos as a proxy for neuronal activation reflects only the final timepoint, after the completion of PR task. Also, in a PR task, unlike in a FR schedule, the response requirement changes over time so mice cannot generally predict how much work is required to obtain a reward based on previous trials. Therefore, although we can assume a similar hierarchical progression of activity within the striatum, the variable nature of the PR task could impact the behavioural and cellular response differently than suggested for the FR framework (Segovia et al., 2012).

However, we conclude that the perseverative and inflexible behaviour in the ephrin-A2/A5^-/-^ mice used in this study (Moretti et al., 2021) is likely not due to habit formation, because neuronal activity in these mice is mostly localised in regions responsible for goal-directed behaviour. However, previous studies have shown other behavioural patterns in ephrin-A knockout mice that suggest compulsive or perseverative tendencies. Self-injurious compulsive grooming behaviour similar to some behaviours seen in autism spectrum disorder has been reported in a different strain of ephrin-A knockout mice and attributed to abnormalities in the corticostriatal pathway (Wurzman et al., 2015) Although we saw repetitive responding pattern in the ephrin-A2/A5^-/-^ mice in our study, we did not observe self-injurious compulsive grooming. Further evidence of perseverative behaviour in ephrin-A2^-/-^ mice comes from maintenance of a responding pattern during a reversal task, suggesting insensitivity to the reward (Arnall et al., 2010). This perseverative behaviour was broadly attributed to abnormalities in the orbitofrontal cortex and the circuitry involving the thalamus, striatum and the prefrontal cortex. Therefore, perturbations in ephrin-A signalling appear to be consistently linked to compulsive and perseverative behaviour, suggesting that although dopamine abnormalities play a key role in such behaviours, it may be that disrupted organization of neural circuits may be an additional mechanism underpinning perseverative and compulsive behavioural patterns. Neurological disorders often have genetic and developmental factors contributing to their presentation (Ting and Feng, 2011). Therefore, future studies with ephrin-A2/A5^-/-^ mice could give insight into how disruption of neural circuit organization during development have behavioural impacts (Pasquale, 2005; North et al., 2013). Additionally, applying LI-rTMS and its history of successful network reorganization in ephrin models (Rodger et al., 2012; Makowiecki et al., 2014; Poh et al., 2022) to more established models of neuropsychiatric disorders, could be an interesting avenue to elucidate therapeutic potential and mechanisms of rTMS.

### 4.2 c-Fos colocalization with MSNs and interneurons in the striatum

MSNs form the major population of striatal neurons (95%) with the remaining 5% being interneurons (Melzer et al., 2017). These interneurons are quite diverse and have been classified into two major classes, cholinergic interneurons and GABAergic interneurons (Kawaguchi et al., 1995). In our study, we saw an increase in active MSNs (identified by c-Fos-DARPP32 colocalization) in DMS of LI-rTMS-stimulated but not sham stimulated ephrin-A2/A5^-/-^ mice. This suggests that LI-rTMS specifically enhances DMS MSN activity, which may contribute to delaying the shift in striatal activity to DLS. If LI-rTMS can potentially delay an accelerated shift to habitual behaviour, it would be interesting to investigate whether higher dose or more prolonged rTMS treatment could reverse the hierarchical recruitment of striatal regions by shifting activity from the DLS towards regions that are involved in early acquisition of a behaviour, perhaps through activation of DMS MSNs. Such an effect would be relevant to reversing DLS activity associated with overreliance on habits as seen in drug seeking behaviours (Willuhn et al., 2012).

Although MSNs account for the majority of striatal neurons, our qualitative analysis across all experimental groups suggests that roughly 60% and 40% of c-Fos single-labelled cells in the NAc and dorsal striatum respectively, were not MSNs. Therefore, as an exploratory analysis we stained for GABAergic and cholinergic interneurons to identify the nature of the unspecified c-Fos single-labelled cells. In all groups, DMS had more c-Fos^+^-GAD67^+^/DARPP32^-^ colocalization and c-Fos^+^-PV-expressing GABAergic interneurons compared to DLS which could mean more interneuron-related inhibitory activity in the DMS. However, we saw no significant changes in GAD67 activity between groups. In NAc core and shell, across all experimental groups we found that less than 2% of the c-Fos^+^ cells that were colocalised with GAD67^+^/DARPP32^-^ cells. Therefore, although there were no major group differences with GABAergic interneuron activity, the modulatory role of GABAergic interneurons is likely most influential in the DMS due to the higher proportion of active neurons that are GABAergic interneurons. Lastly, in our qualitative analysis we did not find any evidence to support activation of cholinergic interneurons in either dorsal or ventral striatum.

As we tested for all neuronal populations in the striatum, the identity of the high number of unspecified c-Fos^+^ cells is still not clear, as together the staining accounted for only around 40% and 50% of c-Fos^+^ cells in the ventral and dorsal striatum, respectively. There is a possibility that these unspecified c-Fos^+^ cells could be glial cells, however, a confirmatory test in the ventral striatum found that most c-Fos^+^ cells were colocalised with neurons (Moretti et al., 2021).

### 4.3 Relevance

Although the small sample size of the study limits our conclusions, the possibility of rTMS delaying habit formation is exciting. It shines light on potential mechanisms of rTMS as a therapy for disorders such as drug addiction and OCD. rTMS is already used to treat OCD, and there is evidence for anti-craving effects for cocaine (Camprodon et al., 2007) and nicotine (Amiaz et al., 2009). The anti-craving effects of rTMS have been thought to be linked to the ability of rTMS to modulate reward circuitry in the brain. For example, rTMS may counteract addiction-related plasticity by increasing dopamine release and dopamine receptor density, as well as modulating glutamatergic receptor expression (Moretti et al., 2020). Ephrin-A2/A5^-/-^ mice have low dopamine levels in the striatum possibly due to reduced dopaminergic innervation (Cooper et al., 2009; Sheleg et al., 2013). Future studies could look at whether rTMS can alter these abnormal dopaminergic innervations in ephrin-A2/A5^-/-^ mice to normalise dopamine levels in the striatum. Linking this to changes in perseverative behaviour in ephrin-A2/A5^-/-^ mice following rTMS could add to evidence that can support rTMS as a therapy for disorders with overreliance on habits.

### 4.4 Conclusion

The present study did not fully support the assumptions made by our previous study (Moretti et al., 2021) in which it was hypothesized that neuronal activity in ephrin-A2/A5^-/-^ mice with repetitive and perseverative responding behaviour would have shifted to the striatal region involved in habit formation. Rather, in this study, it was found that activity remained dominant in regions involved in goal-directed behaviours. However, the relative proportion of c-Fos activity in the striatum slightly shifted towards DLS activation in untreated ephrin-A2/A5^-/-^ mice when compared to other experimental groups. This was in line with our hypothesis that ephrin-A2/A5^-/-^ mice would have greater c-Fos activity in striatal regions associated with habitual behaviour and that LI-rTMS would reduce this difference. As a whole, rTMS seems to partly delay the shift from goal-directed behaviour to habit in ephrin-A2/A5^-/-^ mice, and this could be due to the changes in MSN activity in the DMS. Future studies could perform electrophysiological recordings from the striatum to map any causal relationship. Overall, LI-rTMS shows some promise for delaying the shift from goal-directed behaviour to habit and/or compulsive behaviour, on a cellular level. This delay could be further investigated in more clinically relevant animal models as a mechanism behind clinical rTMS effects seen in disorders like drug addiction and OCD.

## 5 Acknowledgements

We thank Dr. Bhedita Seewoo and Mr. Samuel J Bolland for their guidance on creating the mouse brain atlas for this study.

## 6 Author contributions

**Maitri Tomar:** Methodology, Writing - original draft, Writing – review & editing, Formal analysis, Investigation, Visualization. **Jennifer Rodger.** Writing – review & editing, Supervision. **Jessica Moretti.** Conceptualization, Writing – review & editing, Methodology, Formal analysis, Visualization, Supervision.

